# Mispatterned motile cilia beating causes flow blockage in the epileptic brain

**DOI:** 10.1101/2021.10.29.465023

**Authors:** Regina J. Faubel, Veronica S. Santos Canellas, Jenna Gaesser, Nancy H. Beluk, Tim N. Feinstein, Yong Wang, Maya Yankova, Kalyani Bindu, Stephen M. King, Madhavi K. Ganapathiraju, Cecilia W. Lo

## Abstract

Beating of motile cilia at the brain ventricular surface generates rapid flow in an evolutionary conserved pattern mediating the transport of cerebrospinal fluid, but its functional importance has yet to be demonstrated. Here we show disturbance of this transport may contribute to seizure susceptibility. Mice haploinsufficient for *FoxJ1*, transcription factor regulating motile cilia exhibited cilia-driven flow blockage and increased seizure susceptibility. Mutations in two epilepsy-associated kinases, Cdkl5 and Yes1, in mice resulted in similar cilia-driven flow blockage and increased seizure susceptibility. We showed this arises from disorganized cilia polarity associated with disruption in the highly organized basal body anchoring meshwork. Together these findings suggest mispatterning of cilia-generated flow may contribute to epilepsy and thus might account for seizures unresponsive to current seizure medications.

**One sentence summary:** Epilepsy is associated with disturbance of cilia motion and mispatterning of fluid transport in the brain ventricles.

## Main Text

Homeostatic imbalances in brain physiology underlie neurological disorders such as epilepsy, a condition characterized by recurrent seizure affecting 1% of the population. Standard pharmacological treatment targeting neuronal synapses is ineffective for up to a third of epilepsy patients, suggesting other pathogenic mechanisms (1). The recent identification of several epileptic disorders with cilia related genes, including those that affect motile cilia function (2–4), suggests a role for cilia related defects in seizure, especially those unresponsive to current seizure medications. A role for motile cilia in the regulation of brain homeostasis is further suggested by the finding of hydrocephalus, an excessive accumulation of cerebrospinal fluid (CSF), in mice with mutations disrupting motile cilia function (5).

Motile cilia are microtubule-based organelles that project from the cell surface of specialized epithelia lining the airway and the brain ventricular system. Motile cilia beat synchronously to generate directional flow, mediating clearance of mucus in the airway, and transport of cerebrospinal fluid (CSF) along the ependymal surface of the brain ventricular system (5, 6). We previously characterized this cilia-generated flow in the ventral third ventricle (v3V), a recessed cavity lined by neurosecretory nuclei of the hypothalamus (7). Indicative of their functional relevance, motile cilia form a highly regulated fluid transport system, the cilia-based flow network that is conserved among mammals, and changes predictably by time of day and night with the switching of cilia beating direction by so far unknown mechanisms (*7*).

Here, we investigated the potential role of mispatterned ependymal cilia-mediated fluid transport in increasing seizure susceptibility and the role of several kinases therein, including mice with Cyclin-Dependent-Kinase-Like 5 (CDKL5) deficiency modeling the early onset epilepsy in the CDKL5 deficiency disorder (CDD) (*8, 9*).

## Results

### Cilia-mediated clearance is critical for seizure susceptibility

Cilia-mediated fluid transport along the brain ependymal surface of the ventral third ventricle (v3V) was previously mapped (*7*) in explant preparation (Fig. 1A). Thus placing the v3V in medium containing fluorescent beads, the stereotypic pattern of fluid transport along the ventricular surface was revealed. This is characterized by an inflow stream located at the narrow anterior duct (ad) and an outflow stream at the posterior duct (Fig. 1B), indicating rapid directional transport of substances through v3V. This would predict in vivo circulation of substances along the ependymal surface the brain ventricular system. Computational modelling of fluid dynamics supports the importance of cilia-generated flow for clearance from the v3V, especially in more recessed areas such as the v3V (fig. S1).

**Fig. 1:**
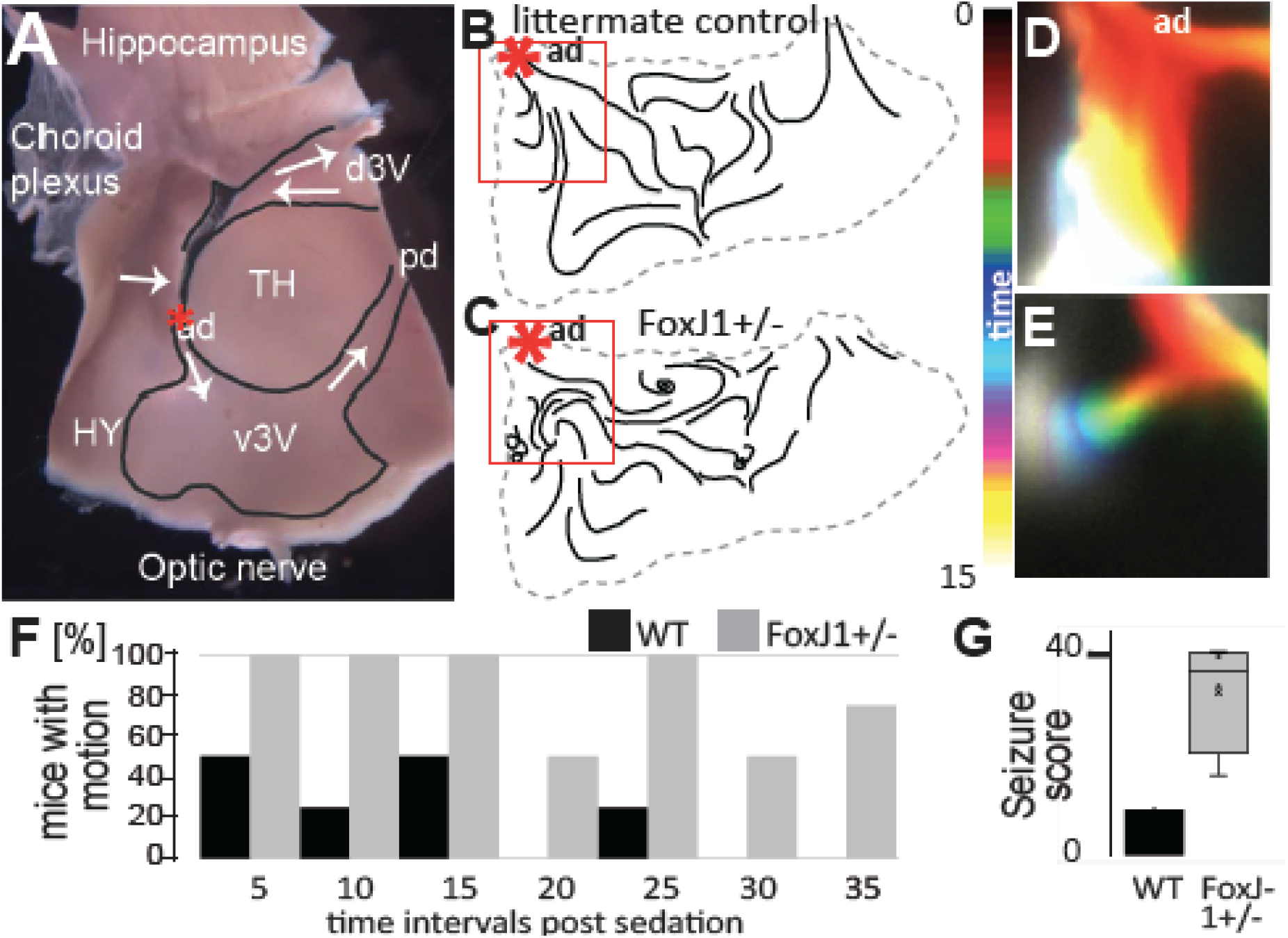
Flow blockage associates with seizure susceptibility. (**A**) Explant with dorsal (d3V) and ventral third ventricle (v3V, black outline) facing the hypothalamus (HY) and Thalamus (TH) and cilia-generated flow (white arrow) via the anterior (ad) and posterior duct (pd). (**B**,**C**) Flow maps show tracing of cilia-generated flow. n=4, age=5-6weeks. (**D**,**E**) Time-resolved color-coding of distribution of FITC-dextran following injection into ad (see star in B,C). (**F)** Percent mice that show any seizure-related activity in 5 second intervals after onset of anesthesia. (**G**) Box plot total seizure scores (t-test p<0.0001).

To ascertain whether motile cilia function is critical for CSF delivery into the v3V via the anterior duct, we investigated v3V explants from mice harboring the *Foxj1* knockout (KO) allele. FoxJ1 is a master transcriptional regulator of motile ciliogenesis (*10*). While homozygous *FoxJ1* KO mice fail to form cilia in the brain ependyma and succumb to hydrocephalus, heterozygous *Foxj1* KO mice (*FoxJ1*^*+/-*^) were grossly normal (*11*) and their ependyma is well ciliated with robust cilia-generated flow (Movie S1). Flow maps of the v3V were grossly similar between the *FoxJ1*^*+/-*^ and wildtype mice, except at the anterior duct where a striking circular flow or whirls were observed in the *FoxJ1*^*+/-*^ mice (Fig. 1C). This was confirmed by tracking FITC-dextran injected into the anterior duct to mimic freshly delivered CSF. While in wildtype mice rapid spread was observed along the v3V surface (Fig. 1D), in the *FoxJ1*^*+/-*^ mice the inflow of injected dye was not observed, indicating blockage of inflow at the anterior duct (Fig. 1E).

To investigate the potential functional impact of this inflow blockage in v3V in the *FoxJ1*^*+/-*^ mice, we used anesthesia induced convulsions as a readout of seizure susceptibility (fig. S2). During the early phase of anesthesia, wildtype mice show occasional twitching of limbs and neck (Movie S2), whereas in the *FoxJ1*^+/-^ mice, more severe twitching of longer duration was noted (Fig. 1F,G), often accompanied by mild to tonic-myoclonic seizures (Movie S3). We note anesthesia induced seizures also can be observed in patients with a history of epilepsy (*12*).

These findings show increased seizure susceptibility associated with blockage of cilia-generated flow, suggesting impaired tissue clearance.

### *Cilia lengthening in patients with CDKL5* deficiency disorder and *Cdkl5 KO* mice

The potential functional importance of ependymal cilia-mediated flow in seizure susceptibility was further investigated by examining the seizure disorder associated with the X-linked gene *CDKL5* referred to as CDKL5 deficiency disorder (CDD). RNAseq and ChIPseq profiling indicated the X-linked gene *CDKL5* is a downstream target of FOXJ1. Its protein serine/threonine kinase activity is among the top ten molecular functions most strongly regulated by FoxJ1 (11). CDKL5 is highly conserved, with its kinase domain showing conservation from man to the green algae *Chlamydomonas* (fig. S3). Mutation in the *Chlamydomonas CDKL5* ortholog *lf5* causes defects in the flagella, motile cilia that mediate swimming behavior (*2*).

We recruited two CDD patients, a female (P1) and male subject (P2) (table S1). Both patients began having seizures within weeks of birth that quickly became intractable to standard antiepileptic medications (Supplementary Material). We conducted nasal scrape to obtain multi-ciliated nasal epithelia as a proxy for the brain ependymal cilia. Immunostaining showed CDKL5 localization in the ciliary axoneme and basal body of the control subject (Fig. 2A), but not in the CDD patient, confirming CDKL5 deficiency (Fig. 2B). High speed videomicroscopy showed robust ciliary motion (Movie S4), with cilia lengthening noted in both patients when compared to healthy control subjects (Fig. 2C). We note lengthening of the flagella has also been observed in *lf5* mutants (*2*).

**Fig. 2:**
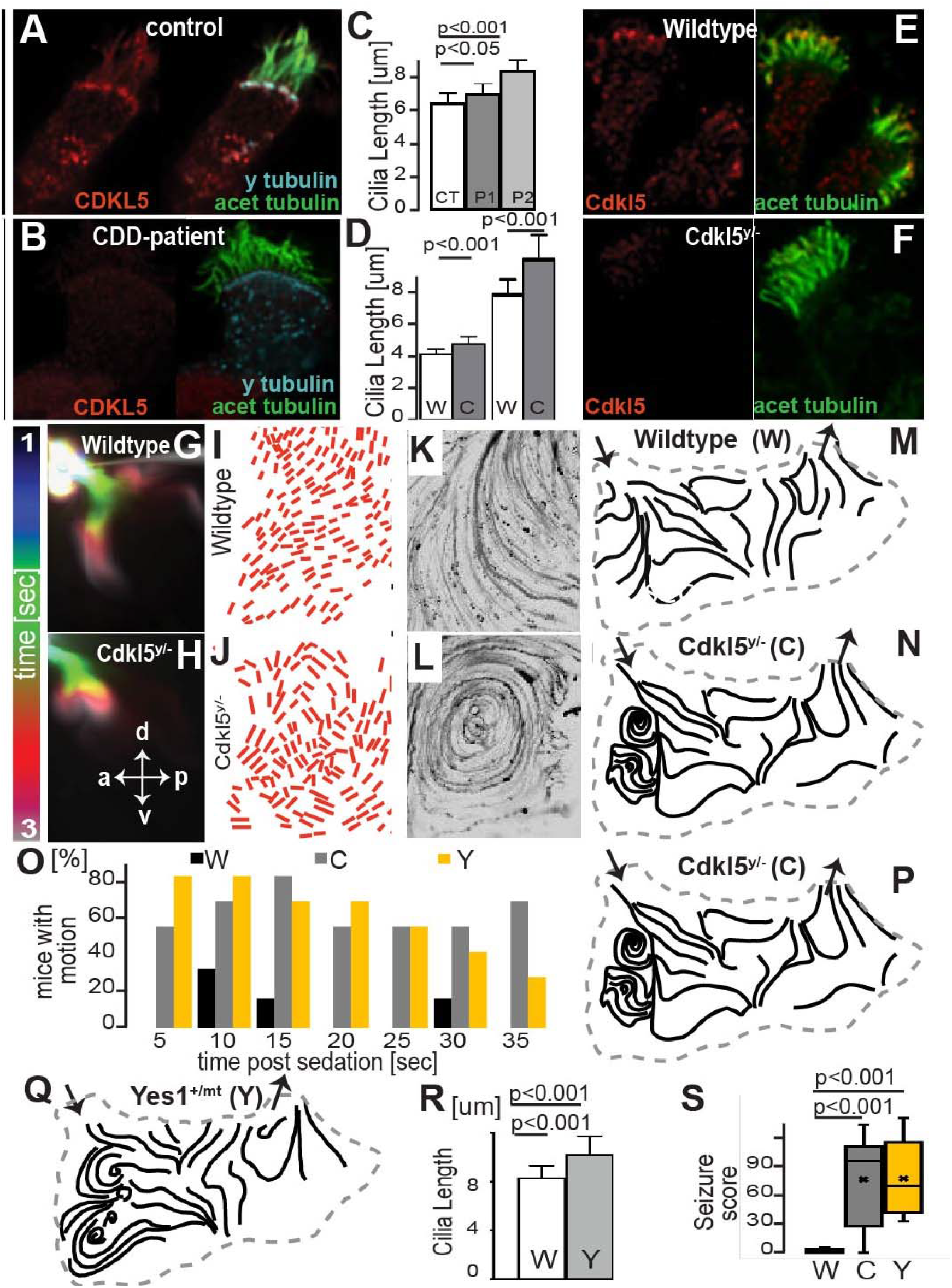
Epilepsy-associated kinases involved in patterning of cilia-generated flow. (**A**) Immunostaining of CDKL5 in human nasal epithelia reveals ciliary localization (**B**) is absent in CDD-patients. (**C**) Motile cilia lengthening is observed in patients and (**D**) respiratory (left) and ependymal cells (right) in *Cdkl5* KO mice (C) vs control (W) mice. (**E**) Cdkl5 staining in motile respiratory cilia (**F**) is absent in *Cdkl5* KO mice. (**G**) Cilia-generated flow from the anterior duct into v3V (**H**) is blocked in *Cdkl5* KO mice (**I**,**J**) with misorientation of cilia beating direction (**K**,**L**) generating circular flow (**M**,**N**) at the a=anterior v3V where fresh CSF enters from the anterior duct. p=posterior, d=dorsal, v=ventral. W: wildtype, C: hemizygous *Cdkl5* KO, Y: Yesl^m/+^ mutant, (**O**) Incidence of seizure like physical motion after anesthesia induction. (**P**,**Q**) Flow map of the v3V after ependyma specific *Cdkl5*-deletion (P) and in the Yes1^+/mt^ mutant mice (Q). (**R**) Ependymal cilia length in Yes1^+/mt^ mice. (**S**) Quantification of convulsions (t-test).

Given many of the pathological findings in CDD patients are replicated in the*Cdkl5* KO mice, including seizures (*8, 9, 14*), we analyzed motile cilia in the tracheal epithelium and brain ependyma of the *Cdkl5* KO mice. Motile cilia in both tissues showed lengthening (Fig. 2D, Movie S5). Antibody staining confirmed Cdkl5 localization in motile cilia of wildtype (Fig. 2E), but not *Cdkl5* KO mice (Fig. 2F). Thus motile cilia in the airway and brain are both subject to regulation by CDKL5, indicating suitability of using the patient nasal epithelia as a proxy for brain ependymal cilia.

### Cdkl5 KO mice show defects in motile cilia and increase in seizure susceptibility

Functional consequence of *Cdkl5* deficiency on ependymal cilia-generated flow was further examined using fluorescein isothiocyanate-(FITC)-dextran to map the pattern of cilia-generated flow in v3V explants of the *Cdkl5* KO mice. While wildtype mice showed the expected directional flow into the v3V via the anterior duct (Fig. 2G), this inflow was blocked in the *Cdkl5* KO mice (Fig. 2H, Movie S6), similar to the previous findings in the *FoxJ1*^*+/-*^ mice. Tracing of cilia motion in this area showed unidirectional cilia beating in wildtype mice, but in the *Cdkl5* KO mice cilia beating polarity was altered (Fig. 2I,J). Further flow mapping with particle tracing using dextran beads revealed a pronounced whirl arising from the disordered cilia beating orientation below the anterior duct (Fig. 2K-N, Movie S7), consistent with whirls also seen in flow maps of the *FoxJ1*^*+/-*^ mice. Similar observation of poorly organized cilia beating direction in the tracheal airway epithelia (fig. S4A,B) points to a role for *Cdkl5* in regulation of motile cilia function overall. Importantly, using the anesthesia induced seizure assay, the *Cdkl5* KO mice were found to have significantly increase in seizures, supporting the role of *Cdkl5* in seizure susceptibility (Fig. 2O). These findings suggest CDD may be a novel ciliopathy involving perturbation of motile cilia function in the brain. However, in contrast to primary ciliary dyskinesia, a ciliopathy involving motile cilia defects in the airway (*5*), electron microscopy showed no evidence of ciliary axonemal defects in either the airway or brain ependymal cilia (fig. S4C,D). However, ciliary membrane protrusions and membrane fusions between cilia were occasionally observed in the *Cdkl5* KO mouse ependyma (fig. S4E,F). Inspection for basal feet showed multiple electron dense masses at each basal body of the ependymal cilia, obscuring their use for determining cilia polarity (fig. S4G,H).

### Cdkl5 deficiency restricted to the brain ependyma causes v3V flow blockage

As *Cdkl5* is expressed not only in the ependyma, but also in neurons and other cell types in the brain, we further investigated whether CDKL5 deficiency in the brain ependyma alone is sufficient to generate the v3V inflow obstruction seen in the *Cdkl5* KO mice. *Cdkl5* deletion to the brain ependyma was carried out using a *Cdkl5* floxed allele (*Cdkl5*^*y/fl*^) in conjunction with a tamoxifen inducible *FOXJ1* promoter driven Cre to target *Cdkl5* deletion to the brain ependyma (FOXJ1-Cre^ERT^) (*15, 16*). Flow maps again showed the same blockage of the v3V inflow in the *Cdkl5*^*y/fl*^ mice carrying the FOXJ1-Cre^ERT^ (Fig. 2P), but not in controls (fig. S5), confirming CDKL5 deficiency in the ependymal epithelium alone is sufficient to replicate the v3V inflow blockage seen in the *Cdkl5* KO mice.

### Ependymal cilia defects and seizure susceptibility in Yes1 Mutant Mice

Given sequence conservation in CDKL5 between *Chlamydomonas* and man is restricted to the kinase domain, this suggests kinase function may be critical to cilia regulation. To investigate this further, we turned to mice harboring missense mutation in the kinase domain of another epilepsy related serine-threonine kinase Yes1 (*17*) (c.T4304C:p.L1435P). *Yes*1 is also a downstream target of FoxJ1 (*13*). Analysis of the v3V from heterozygous *Yes1*^+/mt^ mice showed the same whirl blocking inflow into the v3V (Fig. 2Q). Significantly, airway and ependymal cilia were both lengthened (Fig. 2R), and also observed was increased anesthesia induced seizures (Fig. 2O,S), indicating increased seizure-like activity indicating increased seizure susceptibility. These findings support importance of the serine-threonine kinase pathway in modulating motile cilia and seizure susceptibility.

### Altered ependymal ciliary rootlet meshwork disrupts specification of regional cilia polarity

Cilia length, polarity and coordinated motion are modulated by actin dynamics (*18, 19*), and both Cdkl5 and Yes1 are known to regulate the remodeling of actin (*20, 21*). We conducted confocal microscopy to examine this ciliary rootlet meshwork using antibodies to γ-tubulin to visualize the basal bodies and rootletin antibody to visualize the meshwork. The γ-tubulin stained basal bodies were observed to be embedded in a dense meshwork of rootlet fibers that project to the cell surface, coalescing with similar fibers extending from the meshwork encasing the basal bodies of the adjacent cells (Figure 3A-D). These fibers also appear to ensheath the nuclei of each cell (Figure 3A-D).

**Fig. 3:**
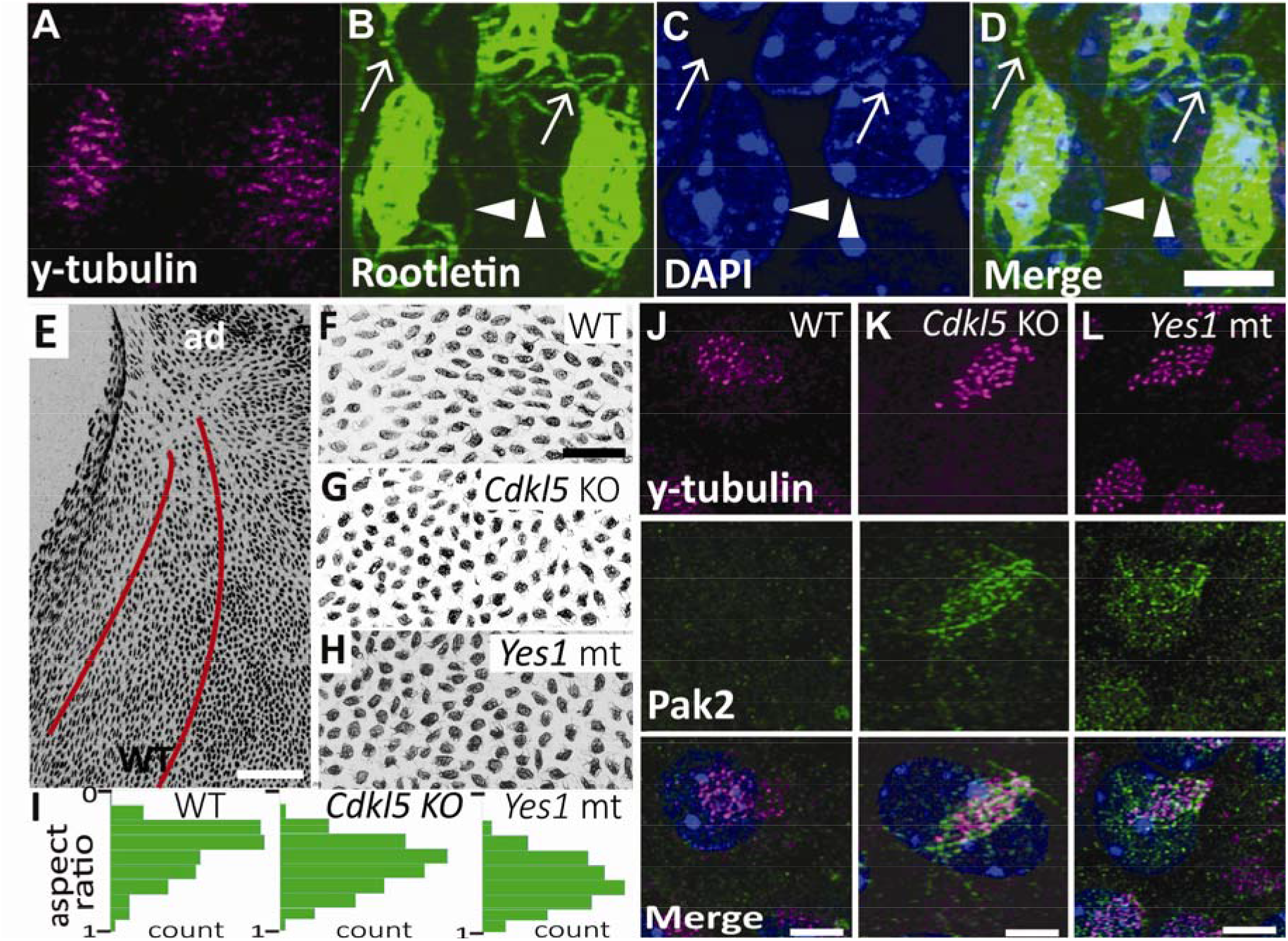
Alteration of cilia anchoring rootlets underlying flow abnormalities. (**A-D**) Immunostaining shows basal bodies are embedded in a meshwork of rootletin-stained fibers that wrap around the nucleus (arrowhead) and interconnect between cells (arrow). Bar=5um. (**E**) Immunostaining for rootletin shows an eccentric structure with the long axis pointing in the direction of cilia-generated flow (indicated by red lines). Bar=50um. (**F-I**) Roundness of the ciliary rootlet containing meshwork in the anterior v3V is increased in *Cdkl5* KO (**G**) and *Yes*^+/mt^ (**H**) mutant mice, as shown in the quantification in panel **(I)**. Bar=30um. (**J-L**) Immunostaining reveals colocalization of Pak2 with fibrous structures above the nucleus (blue) in *Cdkl5* KO and *Yes1*^+/mt^ multiciliated ependymal cells, but not in the ependymal cells of wildtype (WT) mice. Bar=3um.

This meshwork was present in the *Cdkl5* KO and *Yes*^+/mt^ mice, but their regional organization was altered. Rather than exhibiting an eccentric or oval shape with the long axis aligned with the cilia beating direction (Fig. 3E,F,I), this rootlet meshwork in the mutants were rounded, showing little or no directional orientation (Fig. 3G-I). As basal body anchoring to the actin meshwork via ciliary rootlets is mediated by ciliary adhesions in analogy to focal adhesions (*22*), we further examined the distribution of Pak2, a p21 (Cdc42/Rac)-activated kinase that is a downstream effector of *Yes1* (*21, 23*) and also known to modulate focal adhesions (*24*). While little or no Pak2 immunostaining was observed in the v3V ependyma of wildtype mice (Fig. 3J), in both the *Cdkl5* KO and *Yes1*^+/m*t*^ mice, prominent staining was associated with fiber-like meshwork embedding the basal bodies (Fig. 3K,L). Together these findings suggest the organization of the ciliary rootlet meshworks may have a critical role establishing the stereotypic flow pattern in the v3V inlet.

### Kinase pathway regulates synchronization and stability of cilia motion

As the rootlet-associated actin network regulates metachronal cilia beating in Xenopus (*19*), we examined for possible disturbance of ciliary motion in the nasal epithelia of CDD patients. This analysis showed a loss of temporal synchronization of cilia beating in the CDD patient respiratory cilia (Fig. 4A-D). Analysis of the cilia stroke angle was increased from 120° in wildtype mice to 160° in the *Cdkl5* KO and *Yes1*^+/mt^ mice (Fig. 4E). This increase in stroke angle indicating more travel in the ciliary beat provides further evidence of altered basal body anchoring in the meshwork. These findings support a role for the cilia anchoring meshwork in causing regional disruption in the patterning of cilia polarity and cilia mediated flow that could underlie the observed flow blockage in the v3V (Fig. 4F,G).

**Fig. 4:**
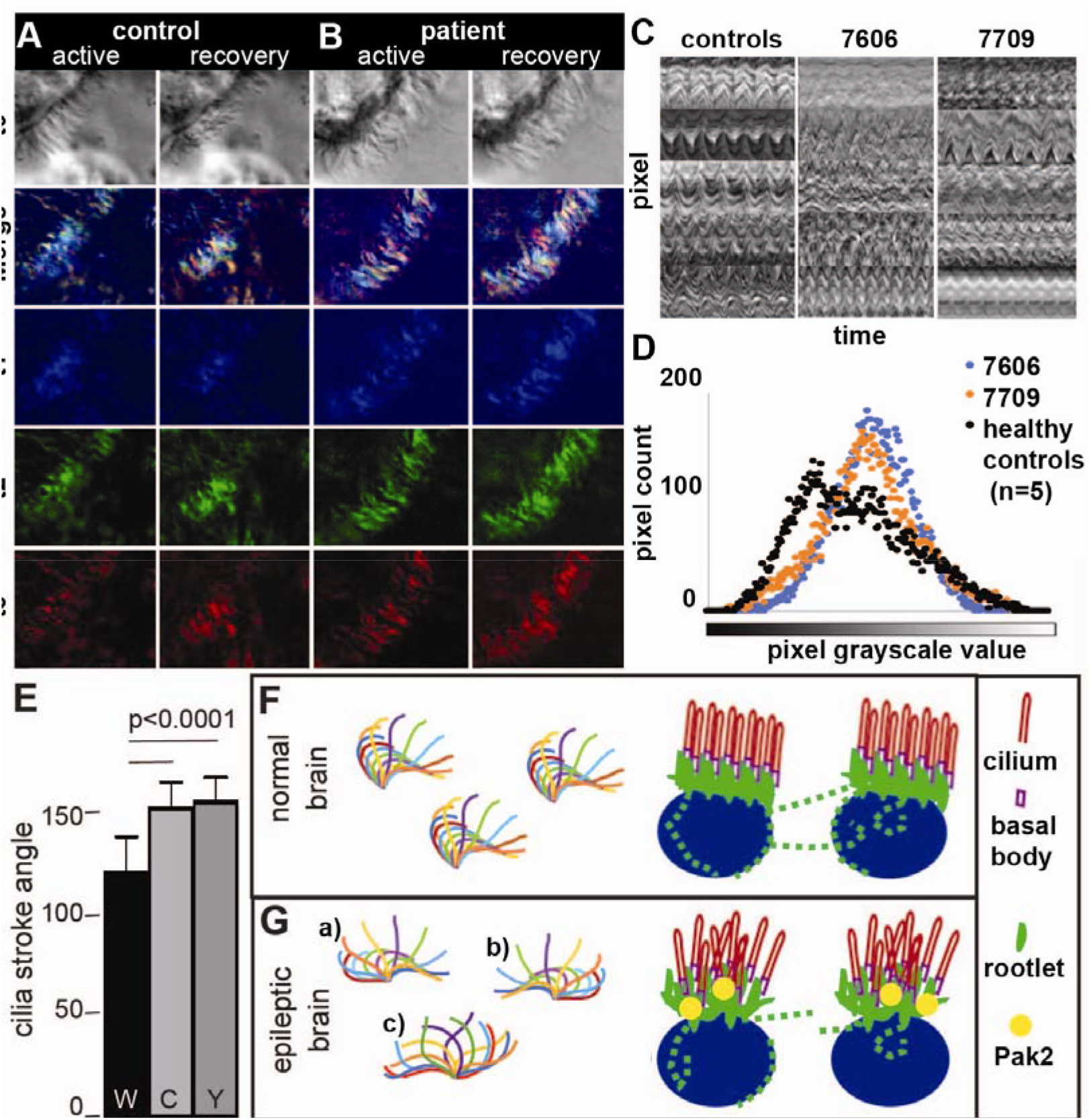
Abnormal ciliary motion associated with altered ciliary rootlet anchoring. **(A**,**B)** Synchronous transitioning of cilia position is visualized in normal human respiratory cells as seen by color-coding three time points during the active and recovery stroke (A). In contrast, respiratory cells from two CDD patients, synchrony was not observed (B). **(C**,**D)** Kymographs were obtained from the ciliary motion (C) and quantification of their grey scale intensity (D) showed a biphasic distribution in the 5 control subjects, while a single peak was observed in the 2 CDD patients. **(E)** The angle of cilia motion is increased in *Cdkl5* KO and *Yes1*^*+/mt*^ mouse ependyma from 120° to 160°. **(F**,**G)** Wildtype mice (F) exhibit coordinated ciliary motion (diagram on left) with well polarized basal body anchoring meshworks (diagram on right), while mutant mice (G) prone to seizures show abnormal ciliary motion (diagram on left) with disrupted rootlet organization (diagram on right). This is associated with ectopic Pak2 recruitment to the meshwork.

### Seizure Related Genes and Pathways

To investigate the possible larger genomic context for seizure susceptibility involving the three kinases, CDKL5, YES1, and PAK2, we constructed a protein-protein interactome (PPI) with these three proteins. PAK2 and YES1 are direct protein-protein interactors, with PAK2 being a downstream effector of YES1 and CDKL5 (fig. S6). The constructed PPI network comprises 172 genes that included the recruitment of five additional cilia proteins (CDC42, NPHP1, GAS8, CETN1, PDCD6IP), four proteins associated with ependymal defects, three of which are cilia proteins(CDC42, BICD2, GAS8, PDCP6IP), and 13 proteins associated with hydrocephalus (CETN1, PTEN, SLC9A3R1, DLG4, PDCD6IP, CASP3, TP53BP2, GFAP, GAB1, SOCS7, BICD2, CDC4) (fig. S6, Data 1). Also recovered were ten proteins with seizure phenotypes in mice (PTEN, TYRO3, DST, GFAP, BRAF, RASA1, FYN, APP, SYN1 and FMR1) (nodes with black dots in fig. S6), and three associated with human epilepsy (nodes with black dot in fig. S6), one of which, FMR1, is also known to cause epilepsy in mice (black outlined node with black dot in fig.S6)

Pathway enrichment analysis of the PPI recovered many relevant pathways such as actin filament organization, small GTPases mediated signal transduction, cell substrate adhesion, establishment of tissue and cell polarity (fig. S6; Data S1). Together findings from the PPI analysis further support CDKL5, YES1 and PAK2 being part of a kinase-network regulating cytoskeletal organization important for the specification of cilia polarity. Of note, focal adhesion kinase (PTK2) is a Yes1-interactor and also a known downstream effector of Pak2 (*25*), supporting the importance of kinase function in modulating ciliary adhesions in orienting cilia in the *Xenopus* epidermis (*22*).

To obtain a broader overview of the mechanistic underpinning for seizure disorders, we retrieved the ∼900 epilepsy-related genes annotated in the Mouse Genome Informatics database for Gene Ontology enrichment analysis (fig. S7). No significant enrichment was found for terms related to neuronal activity, neurotransmitter pathways or for ion channels, consistent with a previous analysis of a smaller group of genes (*1*). Instead, terms significantly enriched were largely related to metabolic function, such as biotin metabolism, urea cycle and metabolism of cofactors. Together with our findings showing the important impact of ependymal ciliary function on seizure susceptibility, this would suggest perturbation in cilia mediated ependymal flow may contribute to epilepsy disorders via the disturbance of CSF-metabolic homeostasis.

## Discussion

Our findings point to a functional link between the patterning of brain ventricular surface flow and seizure susceptibility in the *Cdkl5* KO and *Yes1* mutant mice. The analysis of the *FoxJ1*^*+*/-^ mutant and *Cdkl5*^*y/fl*^ mice carrying the FOXJ1-Cre^ER^ showed this is mediated via a cell autonomous function of the ependymal cilia. We further showed the regional disturbance of flow at the v3V anterior duct that created a whirl blocking cilia-driven flow is associated with disorgnaized cilia polarity. This involved perturbation in the highly interconnected cilia anchoring rootelin-containing meshworks. The homologous meshwork in the *Xenopus* epidermis has been shown to regulate the cellular patterning of basal body spacing and cilia polarity essential for metachronal synchrony of ciliary beating mediating epidermal mucus transport (20). In the v3V ependyma, a similar basal body anchoring meshwork was observed with rootelin fibers projecting towards the cell surface. They appear to interconnect adjacent cells, forming a superstructure interlinking the basal body meshwork. This could provide regional orientation of cilia beating polarity required for directional flow. Consistent with this, in wildtype mice an eccentric orientation was observed aligned with the direction of flow, but this eccentricity was lost in the *Cdkl5* KO and *Yes1* mutant mice. The abnormal recruitment of Pak2 to the meshwork in the *Cdkl5* KO and Yes1 mutant mice would suggest possible disturbance of ciliary adhesions previously shown to mediate basal body anchoring to the actin meshwork. Interestingly, rootelin fibers associated with the ciliary meshwork also were observed to envelop the nucleus, possibly tethering the nucleus to this meshwork superstructure to direct the forces generated by the ciliary beating to provide directional flow.

Based on these findings, we hypothesize the flow obstruction observed in the v3V may disrupt circulation in the brain ventricular system, possibly causing homeostatic imbalance that can contribute to seizure susceptibility. In the context of this model, the rarity of spontaneous seizures in the *Cdkl5* KO mice may reflect diffusion mediated equilibration made possible by the smaller size of the mouse brain. This would predict animal models with larger brains such as pig or sheep might provide a better context to model seizures associated with mispatterning of cilia driven flow, such as observed in CDD. Altogether, these findings point to a crucial role for CDKL5 in conjunction with other kinases in regulating cilia-generated flow that may have critical importance in seizure susceptibility. We conclude the mispatterning of ependymal cilia generated flow may comprise a novel mechanism for epilepsy that may contribute to seizures that are unresponsive to current seizure medications.

## Acknowledgments

We thank Pete Lefebvre for advising about the initial idea that CDD might be a motile ciliopathy. We thank Dr Joe Zhou and Dr Chay Kuo for generously providing *Cdkl5*^fl/fl^ and FOXJ1:Cre mice, and supporting with advice.

## Author contributions

Conceptualization: CWL, RJF

Methodology: CWL, RJF, SMK, YW, MG

Data collection: RJF, MY

Recruitment of CDD-patients: JG, NHB

Data analysis: RJF, VSC, MY

Computational Data analysis: RJF, YW, MG

Visualization: RJF, TNF, YW, MG, CWL

Funding acquisition: CWL, SMK, RJF

Project administration: CWL

Supervision: CWL, RJF

Writing – original draft: CWL, RJF

Writing – review & editing: CWL, RJF, SMK, YW, MG

## Data and materials availability

All data are available in the main text or the supplementary materials.

## Supplementary Materials

Materials and Methods

Supplementary Text

Figs. S1 to S7

Tables S1

References (*1*–*35*)

Movies S1 to S7

Data S1

